# Protein language models can capture protein quaternary state

**DOI:** 10.1101/2023.03.30.534955

**Authors:** Orly Avraham, Tomer Tsaban, Ziv Ben-Aharon, Linoy Tsaban, Ora Schueler-Furman

## Abstract

**Background:** Determining a protein’s quaternary state, *i.e*. how many monomers assemble together to form the functioning unit, is a critical step in protein characterization, and deducing it is not trivial. Many proteins form multimers for their activity, and over 50% are estimated to naturally form homomultimers. Experimental quaternary state determination can be challenging and require extensive work. To complement these efforts, a number of computational tools have been developed for quaternary state prediction, often utilizing experimentally validated structural information. Recently, dramatic advances have been made in the field of deep learning for predicting protein structure and other characteristics. Protein language models that apply computational natural-language models to proteins successfully capture secondary structure, protein cell localization and other characteristics, from a single sequence. Here we hypothesize that information about the protein quaternary state may be contained within protein sequences as well, allowing us to benefit from these novel approaches in the context of quaternary state prediction.

**Results:** We generated embeddings for a large dataset of quaternary state labels, extracted from the curated QSbio dataset. We then trained a model for quaternary state classification and assessed it on a non-overlapping set of distinct folds (ECOD family level). Our model, named QUEEN (QUaternary state prediction using dEEp learNing), performs worse than approaches that include information from solved crystal structures. However, we show that it successfully learned to distinguish multimers from monomers, and that the specific quaternary state is predicted with moderate success, better than a simple model that transfers annotation based on sequence similarity. Our results demonstrate that complex, quaternary state related information is included in these embeddings.

**Conclusions:** QUEEN is the first to investigate the power of embeddings for the prediction of the quaternary state of proteins. As such, it lays out the strength as well as limitations of a sequence-based protein language model approach compared to structure-based approaches. Since it does not require any structural information and is fast, we anticipate that it will be of wide use both for in-depth investigation of specific systems, as well as for studies of large sets of protein sequences. A simple colab implementation is available at: https://colab.research.google.com/github/Orly-A/QUEEN_prediction/blob/main/QUEEN_prediction_notebook.ipynb.

## Background

Proteins are the “working class” of living cells, performing many of the functions critical for life. Learning molecular details about the structure and function of a protein is often a difficult task, as this entails low-throughput and rigorous work. The quaternary state (qs), *i.e*., the number of units assembling together to form a functional unit is an important characteristic of a protein. Many proteins form oligomers to carry out their molecular assignments (1). These oligomers can be of homo- or hetero-oligomeric nature, *i.e*., identical subunits or different subunits, respectively. The oligomeric formation can be obligatory or dynamic (2), and is often crucial for the proper activity of the complex. Examples for oligomeric proteins include the homotetrameric β-galactosidase (3), and the homotrimeric HTRA1 protease (4), where for both the homo-multimer formation is essential for their activity.

Proteins can be classified into families, whether functional (such as GO (5,6) or KEGG (7)), structural (such as ECOD (8), PFAM (9), CATH (10)) or other classifications (e.g. sequence similarity). These classifications are important when comparing proteins to similar members of the relevant cluster. In this context, families of proteins may all adopt the same qs, or alternatively, may contain members that form various qs (Figure 1). This adds a tier of complexity to qs determination, as simply learning or annotation transfer within families will not always yield the correct assignment.

**Figure 1:**
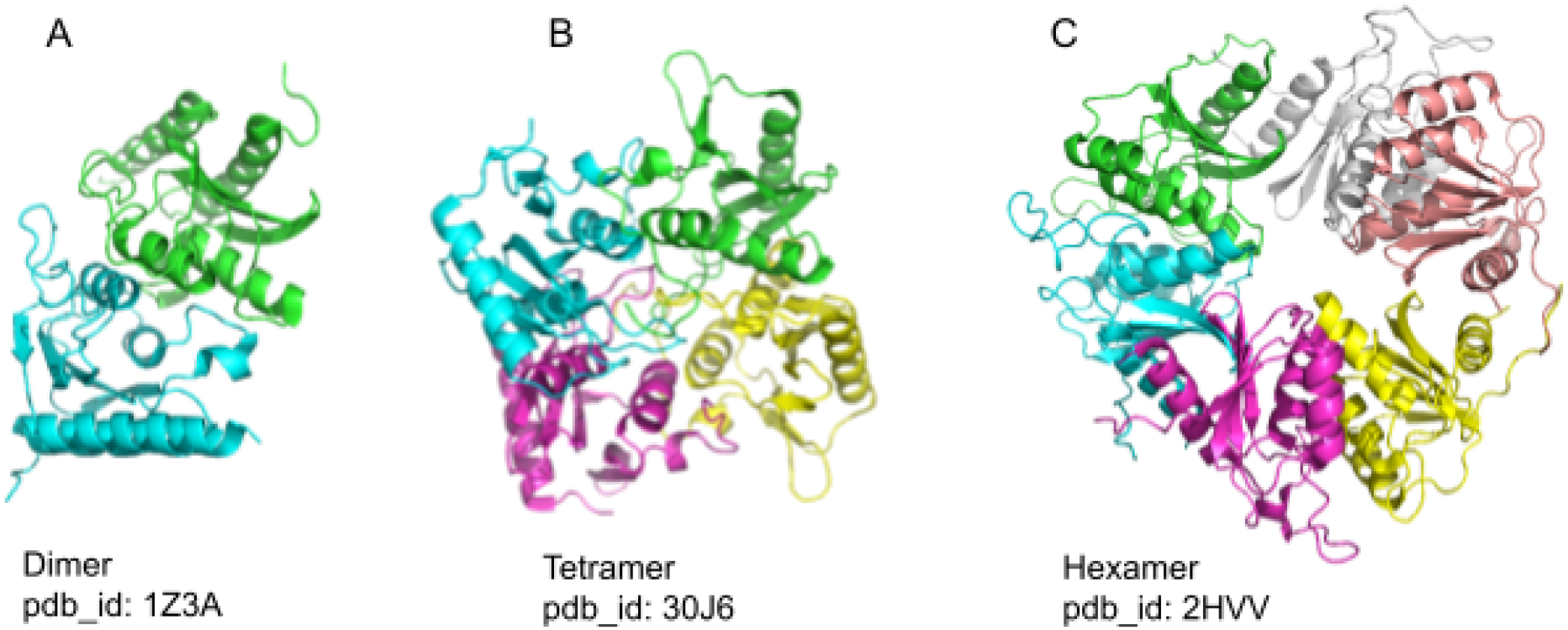
Proteins from ths same fold family can assume different quaternary states (qs). All three proteins belong to the “Cytidineand deoxycytidylate deaminase zinc-binding region’family (ECOD *dCMP_cyt_deam_1*. f_id: 2492.1.1.5), but form either a dimer (A), tetramer (B) or hexamer (C). This demonstrate the complexly of determining qs, as members of the same structural family may form different states.

Usually, experimental approaches are used to determine the qs, such as SEC-MALS, IEX-MALS, ultracentrifugation, to name a few. To alleviate these often expensive and laborious approaches, computational protocols have also been developed for this task, mostly starting from a solved crystal structure (11). As an example, the widely used PISA protocol determines the most likely biologically relevant assembly based on the calculated strength of all different contacts among monomers in a solved structure (12). However, PISA has several drawbacks, the main one being its dependency on information from a solved multimeric structure, where it chooses the correct assembly from within the oligomers in the crystal lattice. Another experiment-based qs predictor, EPPIC takes into account evolutionary information to ascertain the correct qs from a crystal lattice or other solved structures (13). Another tool, GalaxyHomomer, also utilizes solved structures (or generates a model, if no solved structure is available), by selecting the best option among complexes generated by template based docking and *ab-initio* modeling (14). The PROTCID database contains annotations about the quaternary structure of a protein monomer that is based on recurring monomer interactions in distinct solved structures of a protein (15). Finally, QSbio takes into account information about qs states of homologs retrieved from several data repositories (16). This is done by superposing complete complexes and assessing the correct qs by exploiting the available evolutionary information. As for many other structural tasks, AlphaFold has been used in this context as well, where the confidence measures pTM, pLDDT and PAE are used to assess the most probable multimeric state (2,17), or by generating and scoring a variety of complex structures (18).

While it is quite natural to use structural information to elucidate the qs of a protein, there are also disadvantages to the above approaches, in particular, the computationally heavy structure prediction when no experimentally solved structure is available. How far can we then get using only sequence information? A straight-forward way would be to infer qs based on the qs reported for the protein family to which the protein belongs, or from the closest homolog, when available. However, a given protein family may host structurally similar proteins with distinct qs (as, e.g., in Figure 1). How then should the qs be determined from sequence only? More sophisticated sequence representations could help address this challenge.

Learning information about a protein from its sequence is a well established approach in protein annotation, as for example the use of multiple sequence alignments (MSA) to extract evolutionary information (19). Nonetheless, MSA construction is costly in time and computing power, and dependent on the availability of many homologous sequences, and is thus inherently biased towards evolutionary conservation. To address these problems and others, Natural Language Processing (NLP) concepts have been applied to protein sequences, generating protein language models (pLMs) (20–22). Briefly, a Neural Network (NN) is trained on a tremendous amount of protein sequences, learning the connection between the residues, and more specifically, their contextual meaning. Once learned, information can be extracted as embeddings (*i.e*. a numeric vector representation of the protein). These embeddings are subsequently used in a transfer learning step as input for supervised learning. The resulting embeddings are implemented for various tasks, ranging from predicting the protein secondary structure, localization and characteristics, to predicting three dimensional models (22,23). The embeddings are very powerful, which stems in part from the fact that training was carried out when generating the embeddings, therefore the transfer learning step can be performed on a (relatively) small dataset (e.g. (24)).

Training a method for qs classification is challenging, for many reasons, in particular due to the inherently unbalanced data, with monomers outnumbering all other classes (in, e.g., the QSbio dataset; see, Figure 2 below). The small classes have a dozen or even less entries, many of them with high sequence similarity, thus clustering together and providing less additional information. Moreover, the changes needed to shift a sequence from one qs to another may be very small, involving as few as 5 residues and in cases even a single point mutation (4,25).

**Figure 2:**
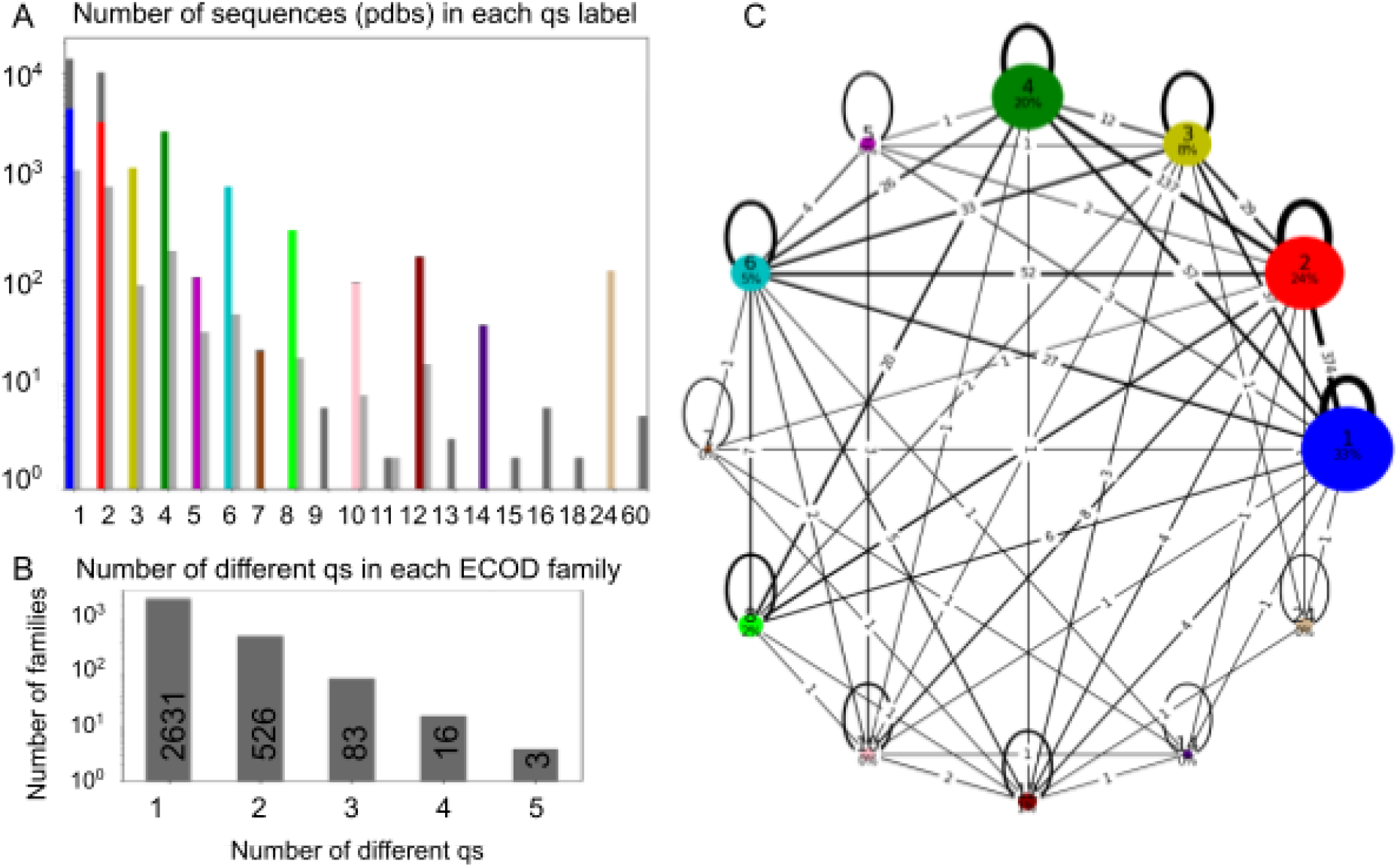
Overall data statistics and distribution of the quaternary state dataset used In this study. **A.** Distribution of qs: For each qs (x-axis) the numberof entries is shown (y-axis, in log-scale). Left and right bars show the training set (in color, after downsampling, dark gray: samples removed far training) and ho?d-out set (light gray), respectively. Downsampling was necessary due to the uneven distribution of the data, which is significantly skewed towards monomers, followed by dimers, trimers, tetramers and hexamers. **B**. Distribution of count of families (y-axis) with different qs states wiihin a given ECOD family (x-axis). While most families show the same qs for all their entries, a significant fraction contains a diverse set of qs (exact numbers are included Inthe boxes). **C.** Details of the composition of qs states in different ECOD families, shown as a network, where the nodes represent different qs (colored as in A.), sized according to their amount (and percentage indicated). The edges represent families containing two different qs slates, with width proportional!to the amount (and numbers Indicated). Note that **B** and C show numbers in the training set, after downsampling and removal of small groups.

In this study we examine the power of pLM embeddings, derived from the pre-trained pLM ESM2 model (23), to capture and consequently classify the qs of proteins. We assemble our training set using curated data from QSbio, including only homomeric proteins, and using only high confidence entries. Our goal in this research is twofold: First, we wish to examine to what extent the embeddings hold the capacity to capture qs, if at all. For this purpose we generate training and test sets, which are available for future research. Second, we built a useful classifier termed QUEEN (QUaternary state prediction using dEEp learNing), which could be employed both for low-throughput manual examination of protein sequences, and especially for high-throughput large scale classifications. One of the major advantages of using pLM is the speed of inference, once the embeddings have been generated. We show that ESM2 embeddings can indeed be used for this task, with performance beyond that of simple sequence homology-based annotation transfer. We build a MultiLayer Perceptron (MLP) model using the embeddings as input, and explore the resulting classifier, its successful predictions and its failures.

## Results

### Organization of the dataset

The QSbio database contains carefully curated information about the multimeric state of different proteins for which the structure has been determined (16). The starting point of our study is a dataset extracted from QSbio, including only homomeric proteins of highest confidence, and only a single annotation per protein data bank (pdb (26)) entry (a total of 31,994 unique protein sequences, see Methods for full details). In this redundant dataset, each unique protein sequence is included as a separate entry, where very similar sequences will most often, but not always, have the same qs annotation. We separated this dataset into a training and a validation (hold-out) set, so that sequences with over 30% sequence identity would be in the same set. We then used the structural domain-based ECOD database (8) to cluster the sequences into domain families (at the family structural similarity level, “f_id”) for their further investigation. Overall, this dataset covers 19 different qs, ranging from monomers to 60mer homopolymers (Figure 2A), and many ECOD families contain representatives of various qs (Figures 2B and 2C, see also Supplementary Figure S1 and Supplementary Tables I & II for detailed information about the dataset and results).

### Distinction of different as states by the language model

First, we generated embeddings for each entry in the database. Each protein is represented by one embedding vector, which is obtained as the average of the vectors of the different residues in the protein sequence (see Methods for more details). In order to assess the capacity of pLMs to capture qs, we used supervised dimensionality reduction to visually demonstrate that the data indeed clusters by qs (Figure 3). The large groups of labels (namely monomers, dimers, trimers, tetramers and hexamers), as well as some other qs are well separated on this map, demonstrating the model’s ability to characterize distinct features of each group. This suggests that the model could be used to predict the qs.

**Figure 3:**
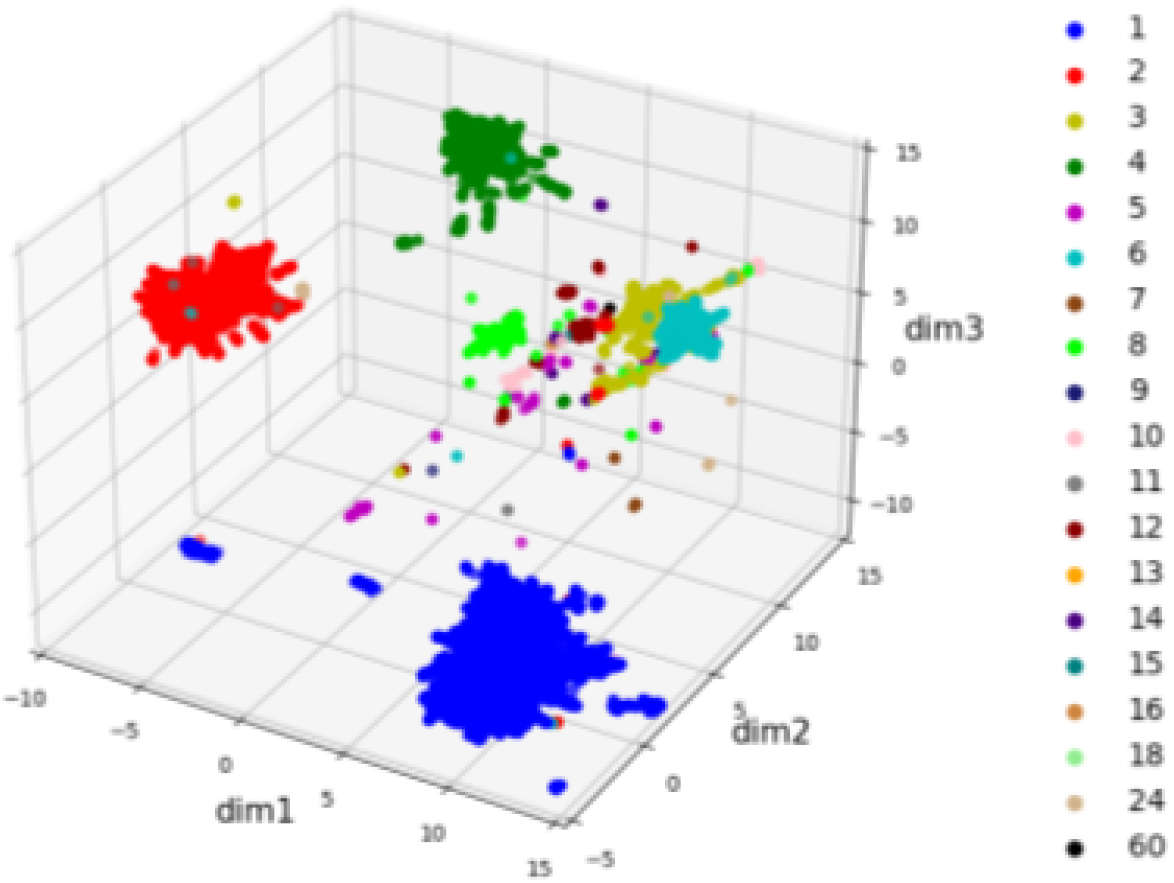
Capture of different qs by the pLM embeddings. Supervised dimensionality reduction by UMAP is able to separate proteins of different qs, in particular for the qs represented by many entries in the dataset, with nice separation for octa mers as well (light green), qs are colored as in Figure 1A. Note that this map includes all the data, *i.e*., both the training and hold out sets.

### Inferring qs by annotation transfer from similar proteins

To assess the ability of QUEEN to correctly predict the qs of a protein, we used annotation transfer to assign a qs to each sequence in the test set (Figure 4A,B and Supplementary Figure S2). In this approach, the qs is inferred from the most similar protein with available annotation, *i.e*. the nearest neighbor in the embedding space (calculated as cosine similarity, see Methods), similarly to previous studies (27). This can be compared to a corresponding annotation transfer based on sequence similarity. Using the embedding space has advantages: a distance can be calculated between any two vectors even if they are very different, since they represent a feature of a protein sequence. This is in contrast to sequence representation that is residue-based, and therefore necessitates sequence alignment, which can be challenging when comparing distant sequences.

**Figure 4:**
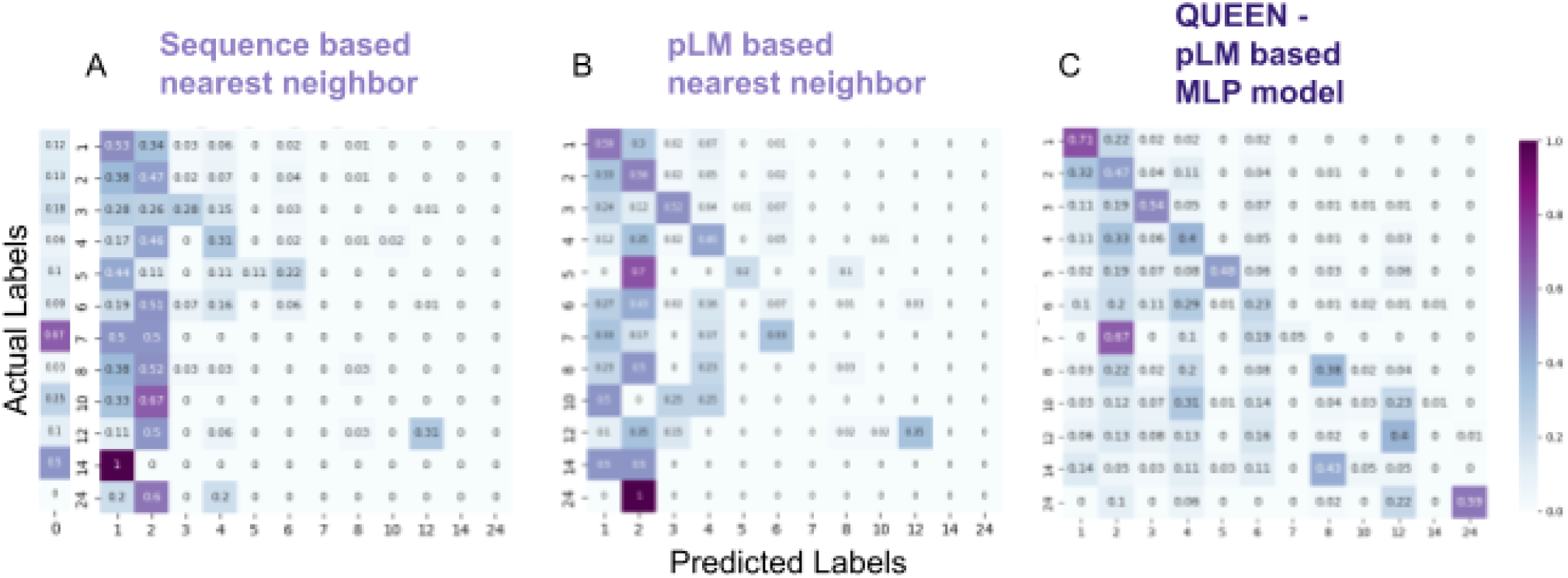
Accuracy of quaternary state prediction by different approaches. The prediction is based on transfer of the qs annotation to each sequence in the test set based on **A**. the closest sequence in the train set (ss determined by blast). The 0-pred?cted column indicates the fraction without any significant blast hit, and consequently no prediction: **B**. the highest similarity in embedding space in the train set *(i.e*. cosine similarity between embedded vectors); and **C**. oQUEEN - a deep learning model trained on the embeddings (see Text). The confusion matrix includes the frequency of cells representing predicted vs. actual label (on x and y-axes. respectively), where a matrix occupying only the diagonal represents full success, while off-diagonal values represent wrong predictions. The balanced accuracy increases from left to right as indicated by the darker diagonal, highlighting improved prediction when moving from sequence, to language model representation, to QUEEN, the MLP model. Results are shown for the test set based on information learned from an independent training set. For corresponding confusion matrices on a redundant set containing also information of sequence similar proteins, see Supplementary Figure S2

Annotation transfer using embedding distance outperformed corresponding annotation transfer based on sequence identity in predicting qs (Table I). This applies to prediction with available prior knowledge (*i.e*., on a redundant set that contains sequence-similar proteins, Figure S2). Importantly, this holds also when prior knowledge is *not* available (*i.e*., for the test set that does not contain any entry with significant sequence identity to the training set): The balanced accuracy increases from 0.15 to 0.23 (Table I, compare Figures 4A,B).

**Table 1:** Performance of different models for the prediction of quaternary states (qs)

|  |  | Annotation transfer, based on |  | pLM MLP model, based on |  |
| --- | --- | --- | --- | --- | --- |
|  |  | Sequence | pLM | ESM embeddings (QUEEN) | Protbert embeddings |
| No information about sequence homologs available <sup>1</sup> | BA<sup>2</sup> | 0.15 | 0.23 | 0.36 | 0.19 |
|  | F1<sup>3</sup> | 0.43 | 0.54 | 0.52 | 0.41 |
| Full homology information available | BA | 0.6 | 0.67 | – | – |
|  | F1 | 0.79 | 0.85 | – | – |

<sup>1</sup> no sequence with >30% sequence available to transfer from; <sup>2</sup>BA: balanced accuracy; <sup>3</sup> F1: F1 score; See Methods for definitions

When the qs of a new sequence is predicted, it is often homologous to previously annotated sequences, which improves prediction (compare Figure S2 and Figure 4, and see Table I). In such a setting, *(i.e*., when including qs information of homologs), we observe a significant separation between cosine similarities used for transfer of correct and incorrect predictions (Figure 5A; with the exception of qs=7; Individual p-values are summarized in Supplementary Table III). Thus, in these cases the cosine similarity can be used to assess whether simple annotation transfer may be sufficient to determine the qs. This is however not applicable for qs assignment without information from homolog proteins, as apparent in Figure 5B.

**Figure 5:**
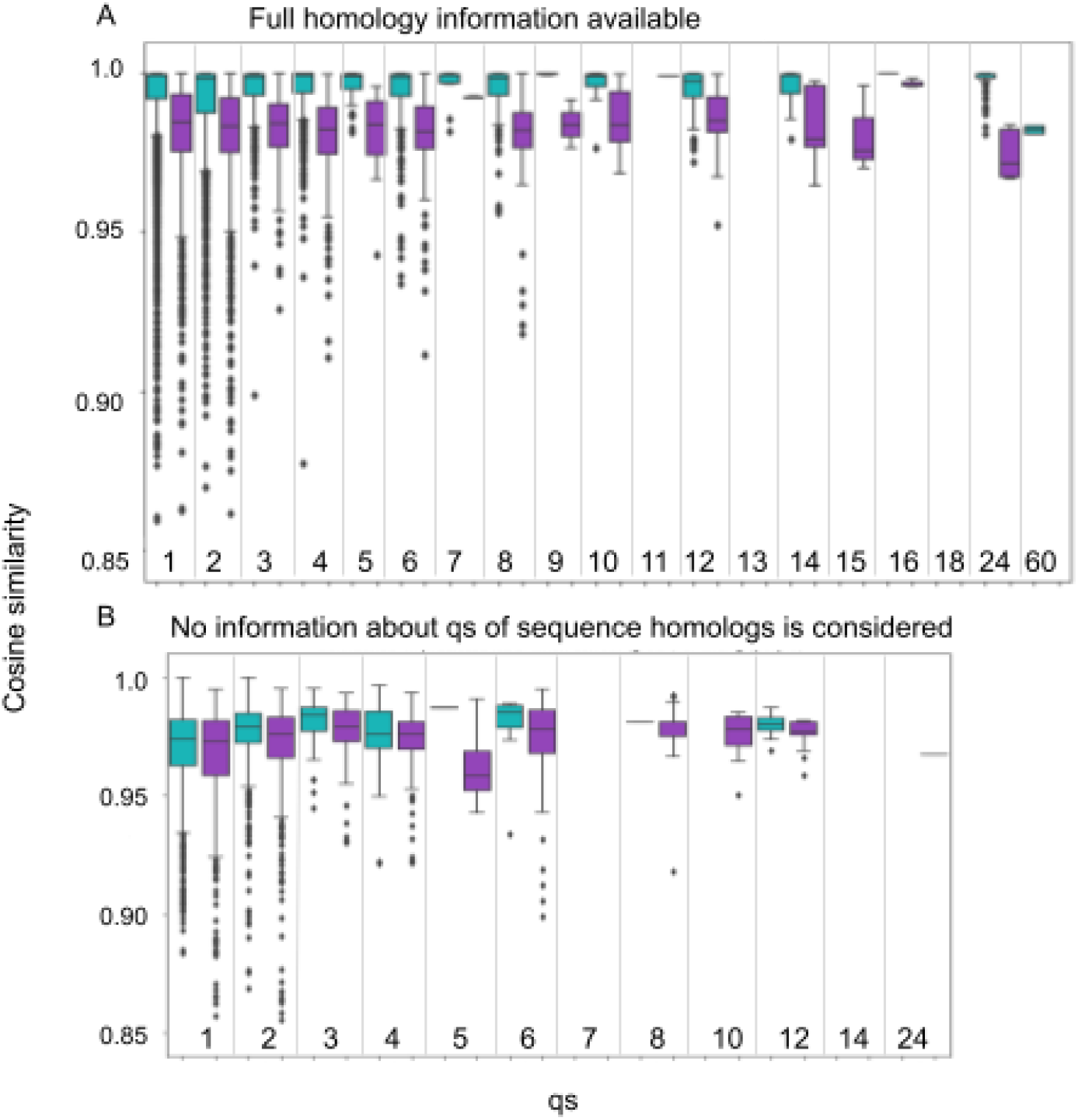
Cosine similarity provide an estimate of the reliability of qs predictions based on annotation transfer from homolog proteins. **A**. Cosine similarity using information that includes qs of homologous sequences. For each qs (x-axis) a box plot depicts the distribution of the cosine similarities for correct (green) and incorrect (purple) qs predictions (y-axis). The plot is capped st 0.85, with *3* dots removed for clarify The two distributions differ significantly (one sided Wilcoxon test, p-values <0,05, except for qs=7, p-value=0.051; see Supplemenlary Table 1). This separation demonstrates the power of embeddings to capture qs, and can be used to assess the confidence of annotation transfers. **B**. Corresponding plot of cosine similarities when information of qs of homologous sequences is NOT included (*i.e*. noentry wilh >30% sequence identity is considered for annotation transfer). inthis case, the two distributions show no significant difference.

### Training a model for qs prediction based on embeddings

The above initial analysis suggests that the embeddings include information about qs. To optimally leverage their use, we trained an embedding-based MLP model to predict qs. We trained and performed hyperparameter tuning on 5 sets of cross-validation within our training set (each time training on 80% to predict a different 20% set, see Methods) to define the final model and its parameters (Figure 4C; Importantly, the hold-out set was not used at any step of training and validation of QUEEN). To achieve improved performance we experimented with several strategies, which highlighted the importance of downsampling of monomers and dimers to obtain a more equilibrated dataset for training (resulting in similar size of monomer, dimer and tetramer qs (see Figures 2A,C). Even after downsampling, the resulting confusion matrix (Figure 4C) still highlights the relatively high success rate of the monomer qs predictions. As for multimers, for many of the wrong predictions, dimers and tetramers are chosen, more so than monomers (see columns of dimer and tetramer predictions in Figure 4C). This hints that QUEEN may have learned to detect multimerization rather than the exact qs, as shown in the striking example of the predominant classification of heptamers as dimers (see also Figure 7 below).

**Figure 6:**
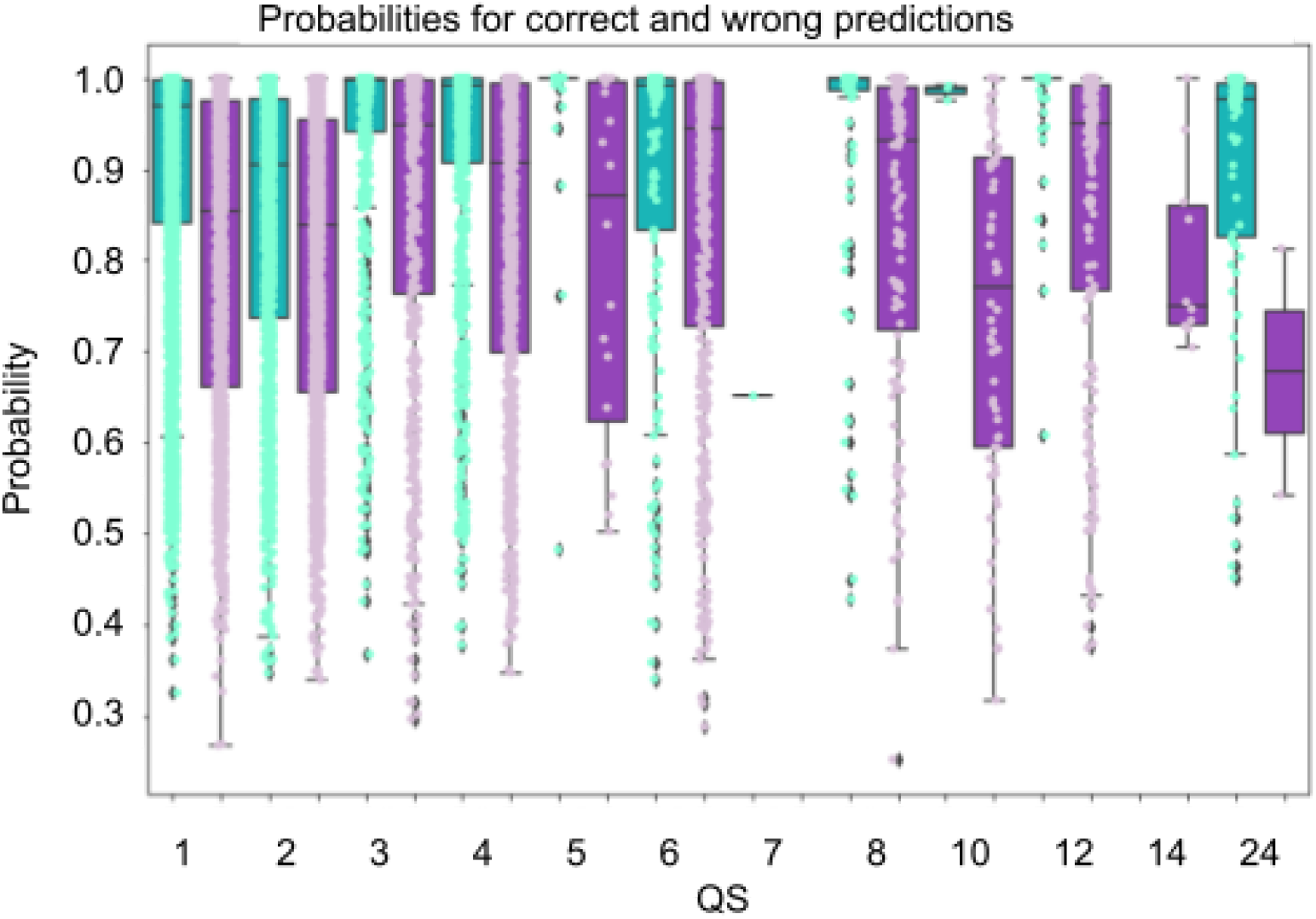
QUEEN predictions shows consistently higher confidence for correct assignments. For each qs(x-axis) box plots and dots show the distribution of the probabilities of correct (green) and incorrect (purple) predictions. Correct predictions are consistently predicted with higher probability than incorrect predictions, for all qs (one sided Wilcoxon test, p-value <0.05). This separation can be used to assess correct predictions, also in the absence of homologs.

**Figure 7:**
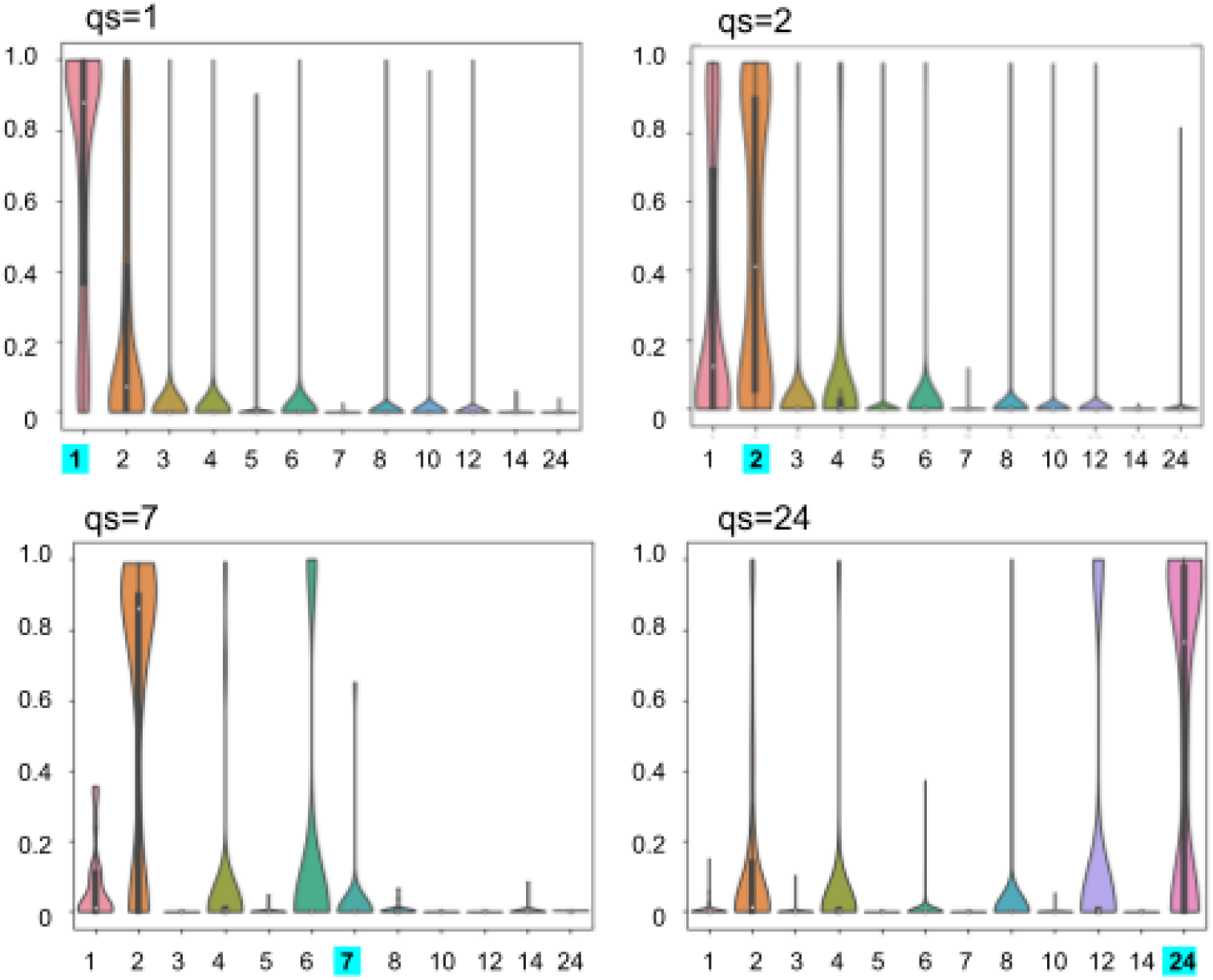
Distribution of predicted probability (‘confidence’) for representative qs categories. The plots show violin plots of the prediction probabilities (y-axis) for different qs labels (x-axis), for actual qs=1,2,7 and 24 (indicated in tho title and highlighted in cyan on the x-axis)

Compared to nearest neighbor annotation transfer, QUEEN shows significant improvement (compare Figure 4C to 4B; balanced accuracy 0.36 *vs*. 0.23, see Table I). While success is not uniform across all labels, correct prediction (> 40% in the diagonal of Figure 4C) is achieved in 7 out of the 12 trained and predicted qs classes.

### Confidence of prediction as indication of success of QUEEN

While QUEEN provides prediction of a specific qs, it is also interesting to examine the underlying probabilities that drive the label prediction and to assess whether they are indicative of successful *vs*. wrong predictions. Reassuringly, comparison of the distributions of the corresponding probabilities (*i.e*. true-positives and false-positives) reveals a significant difference between the two (Figure 6), with p-values ranging from 0.04 to 6*10^-88^ (Supplementary Table IV). Of note, in contrast to the pLM-based annotation transfer for which we can only provide a confidence estimate when information about the qs of homologous sequences is available (based on cosine similarity, Figure 5), using the QUEEN MLP model, this is now also possible for qs predictions that are not based on homolog information (based on the predicted probability).

The distributions of predicted qs states shown in Figure 7 reveal a clear preference for monomers for proteins that form monomers, as expected from the successful prediction for the qs of monomeric proteins (See Supplementary Figure 3 for full plots). In contrast, the distribution of qs predictions for heptamers (that QUEEN did not learn well, see Figure 4C), is much more varied. However, as also apparent in the confusion matrix (Figure 4C), the false negative heptamers are predicted to form different multimerization states, despite monomers being the largest class. This reinforces the notion that protein sequences hold information about multimerization, even though not always about the correct state. 24-mer is another interesting example, where besides high probability of 24-mers, the model also predicts the divisors of 24, with enrichment of the rare 8- and 12-mer classes.

### Examination of diverse families

Incorrect predictions shown in Figure 7 could be a consequence of inaccurate learning for ECOD families that adopt more than one qs. We therefore compared performance for ECOD families for which all members adopt the same qs, to those that adopt diverse qs (families with single members were treated separately, as from one annotated sequence it cannot be concluded whether proteins in such a family can adopt different qs). Overall, the variation of qs within the family *(i.e*., how many different qs it includes) is well captured by QUEEN (Figure 8A). For homogenous families QUEEN predominantly predicts a single qs, while for families with more qs, the number of predicted different qs predominantly corresponds to the actual number of different qs. In this context, performance tends to be slightly improved for single qs families (Figure 8B,C). Moreover, for families with diverse qs, predictions for proteins that adopt the dominant qs of that family are more accurate than corresponding predictions for proteins adopting outlier qs.

**Figure 8:**
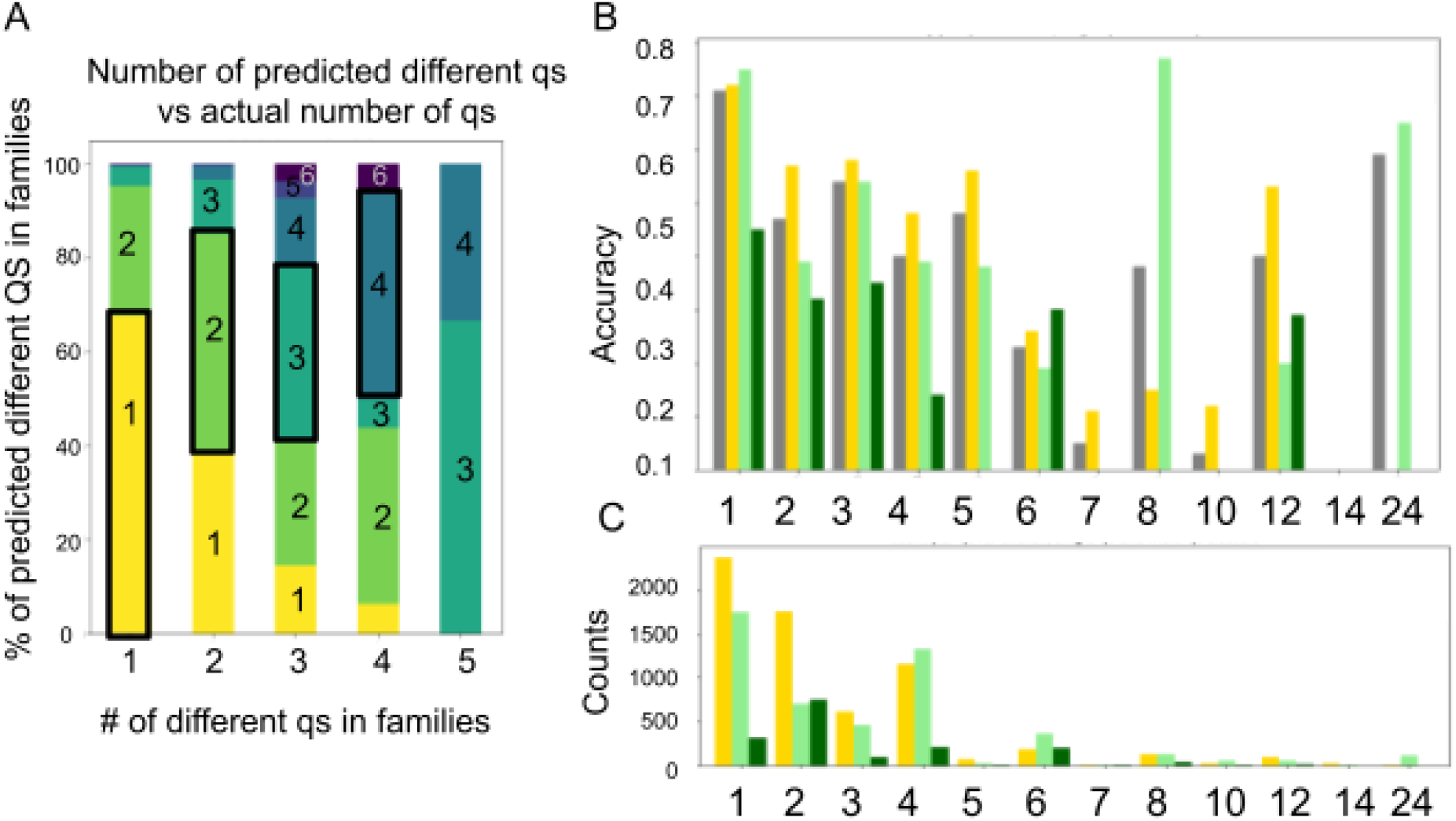
Prediction success and diversity of families with multiple qs. **A**. QUEEN has an accurate representation of lhe degree of diversity of different families: Clusters with multiple qs are predicted as such Shown are the percentage of families predicted to have different qs (y-axis) lor a given number of distinct qc in a family (x-ax?s). Numbers of qe are indicated Inthe boxes and color-coded. **B**. Accuracy (y-axis) for different qs (x-axis). The different bars show four distinct groups wilhin each qs: Overall accuracy (grey), families adopting a single qs (yellow), families adopting multiple qs - predominant qs (light green) and rare qs (dart green). **C**: Corresponding counts of the groups presented in B.

### Comparison of QUEEN performance to other related approaches

How well does this approach perform compared to other related methods? Comparison may provide new insight into the features that determine qs. Of note, to our knowledge, the present model is the first to use sequence information only, in contrast to other methods that all use a solved or predicted structure as input for qs determination or prediction. Not surprisingly therefore, PISA outperforms QUEEN for every class, and EPPIC shows a very similar trend (Note however the better performance of QUEEN for 24mers; Figure 9A). Nevertheless, examination of the degree of complementarity of these approaches suggests that a significant fraction of proteins is correctly predicted only by QUEEN (5-20% for different qs; Figures 9B and C). Therefore, a combination of the different approaches may be beneficial, pending our ability to identify the correct predictions.

**Figure 9:**
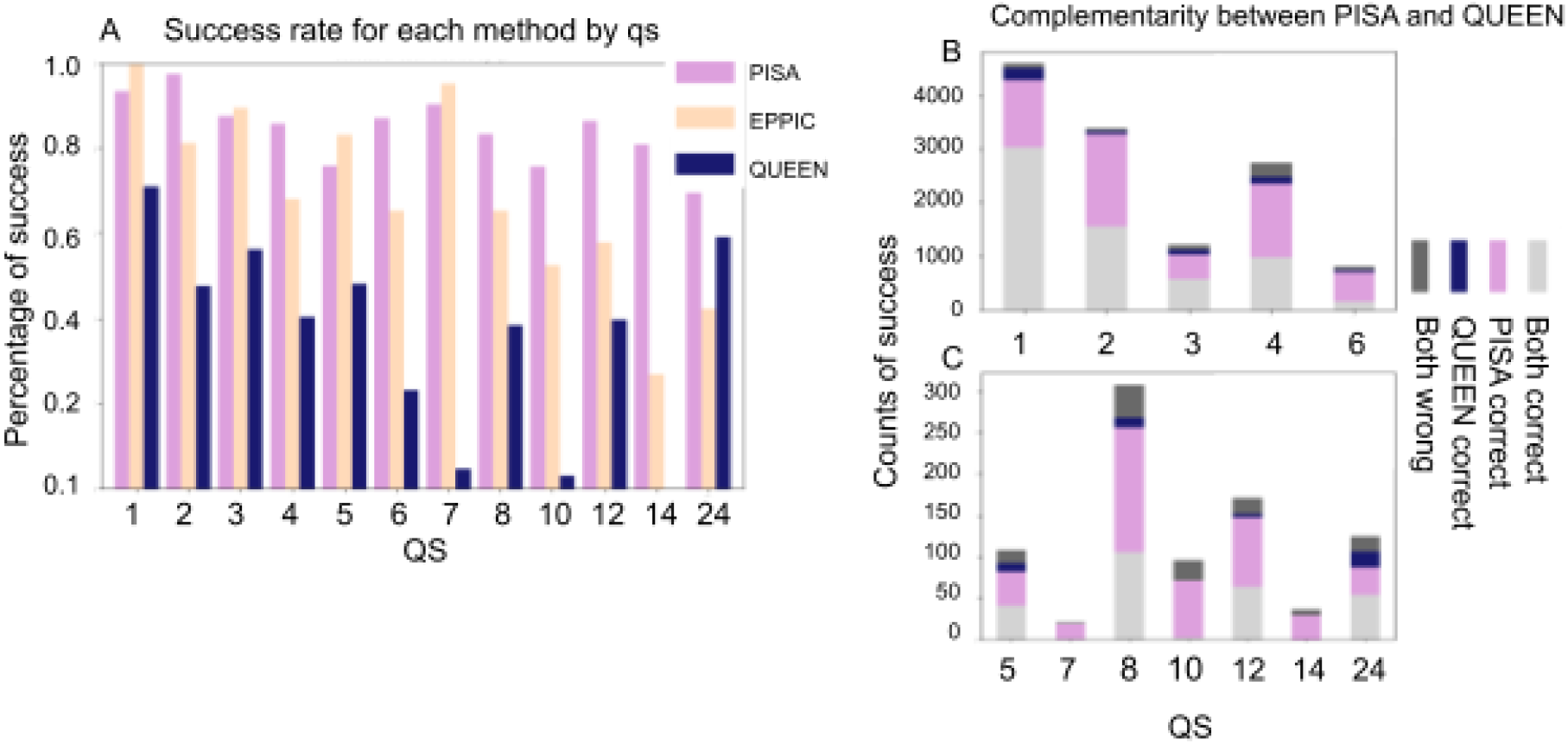
Comparison of success of qs predictions of different methods: Structure-based PISA, EP PIC, and sequence based QUEEN. **A**. Success fy-axis) for different qs (x-axis) for for different Hpproaches. PISA and EPPIC predominantly outperform QUEEN. Note that 1he PISA and EPPIC use not only the structure bu1 also the full crysial Information, and they only choose from tne available options within the crystal lattice. **B and C**. Success (y-axis) colored according to performance by PISA and QUEEN for different qs, highlighting fha significant ovattap, but also the additional potential contribution of QUEEN. The division between B and C la tor danly of the y-axls scale, B shewing tie larger classes 1-4, 9, and C showing the smaller desses 5. 7-24.

### Comparison to other language models - protbert

We show here classification and qs assignment based on embeddings of the protein sequences. These embeddings were generated using the language model developed by Meta - ESM2. We examined whether the source of the embeddings has an impact on the final results, and indeed it does. Repeating this protocol using embeddings from protbert_BFD resulted in worse performance, showing less improvement compared to simple sequence based nearest neighbor annotation transfer (see Table I).

### Robustness of QUEEN performance demonstrated on the independent holdout set

The results presented until now were calculated on the test sets of the 5 cross validation runs. Based on these results, we used the optimized hyperparameter set to define our final model. To evaluate its performance, we now opened the hold out set that we had set aside at the beginning of our study. The performance using sequence or pLM-based annotation transfer on this (smaller) hold out set is slightly better (see Figure 10 for confusion matrices and Table II; compare to Figure 3 and Table I above). Reassuringly, QUEEN performs similarly to what we report for the cross-validation performance, with balanced accuracy slightly lower at 0.3.

**Figure 10:**
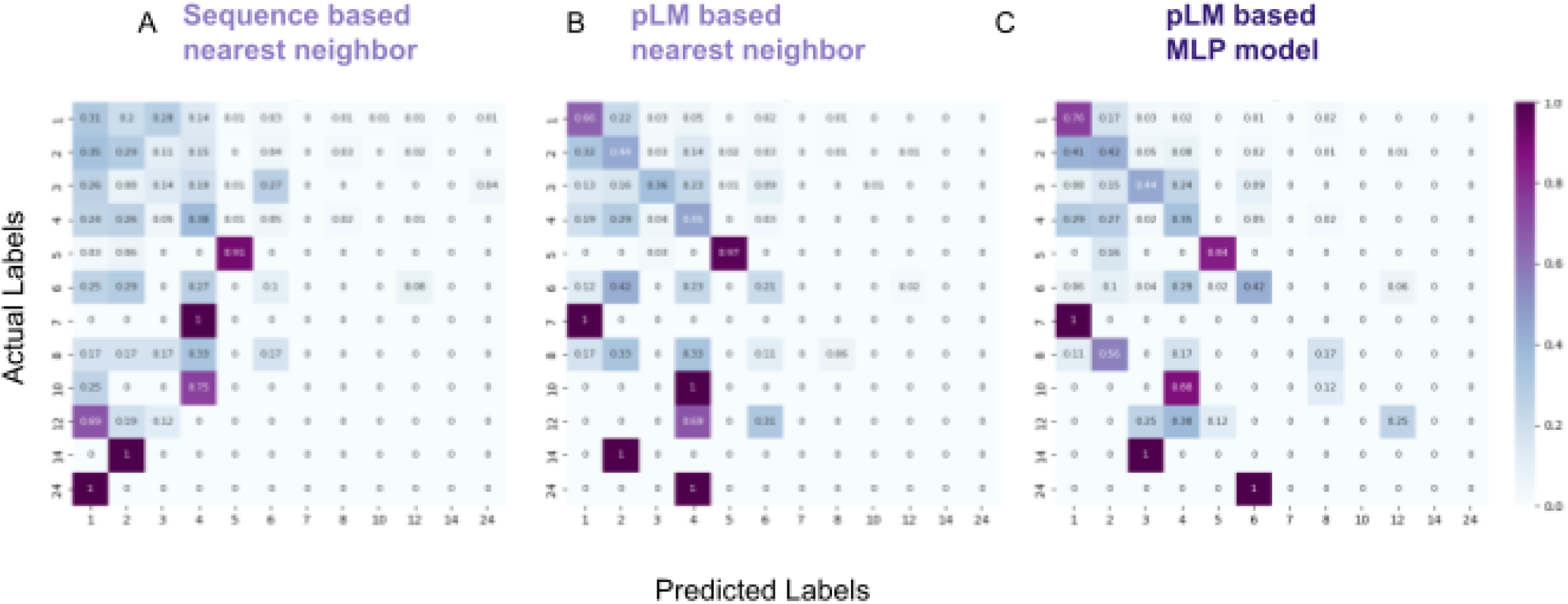
Accuracy of qs prediction by different approaches for the holdout set: The prediction Is based on transfer of the qs annotation to each sequence in the holdout set based on **A**. the closest sequence in the training set (as determined by blast): **B**. The highest similarity in embendding space in tha Training sat (i.e. cosine similarity between embedded vectors); and **C**. QUEEN trained on the embeddkigs (see Text), The confusion matrix includes the frequency of cells representing predicted vs. actual label (on x and y-a*es, respectively), where a matrix occupying only the diagonal represents, full success, while off-diagonal values represent wrong pradicliofis. The balanced accuracy increases from left to right ae indicated by lhe darker diagonal, highlighting improved prediction when moving from sequence, to language model representation, to the deep learning model.

**Table II:**
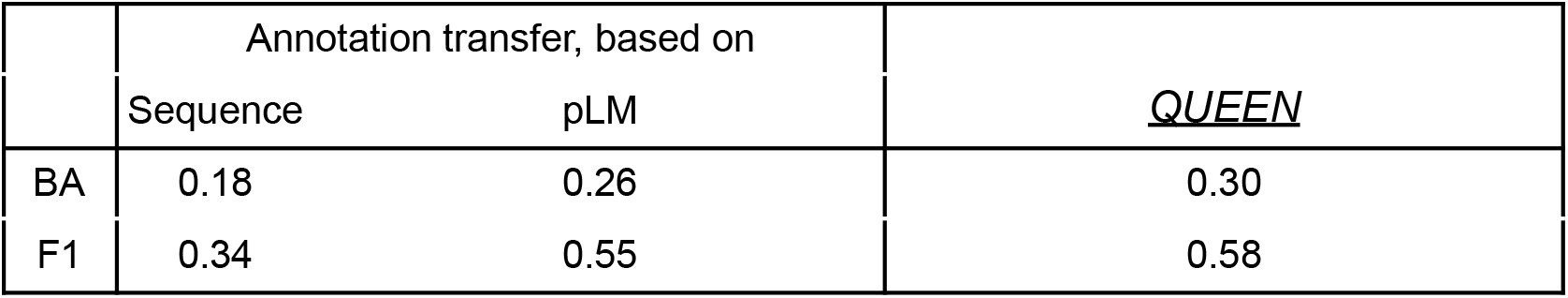
Performance on the holdout set for the prediction of quaternary states (qs)

## Discussion

Protein language models have proven to be most useful to the study of proteins, in particular due to the rich information that their embeddings can provide (28,29). In this work we explore the ability of protein embeddings to capture and classify quaternary states, qs. Our motivation is twofold: First, this allows us to understand how well information about the qs is represented in a sophisticated, yet sequence-based model that does not explicitly consider the protein structure as input. Second, we wished to provide a simple and fast tool to the public for the prediction of the qs of a protein based on its sequence only, to allow for its wide application and hopefully the acceleration of protein research.

Our analysis shows that QUEEN performs better than a sequence-based annotation transfer approach, as well as the use of pLM embeddings only, demonstrating that improved representation of sequence features, and additional learning can improve the representation of qs. Our analysis also demonstrates the difference in performance using different pLMs, as it is better than a corresponding prottrans-based pLM. Overall, QUEEN shows a good ability to distinguish between monomers and multimers. For seven qs success rate tops 40%, a performance which can likely be further improved. Future attempts to predict qs using pLM can profit from including structural information, such as features from a solved or predicted structure, or the use of structural embeddings. As an example, a recent study aimed at prediction of protein function has generated a model that learns PFAM domains, and uses these to infer functional annotation (30). Given information on qs of a larger set of proteins, a similar approach could be implemented for qs prediction.

QUEEN has a “sense” of diversity, *i.e*. sequences from the same family, despite having a similar sequence and similar fold, are not always classified with the same qs. In particular, the degree of variation of qs in different families is overall nicely captured by the model. Therefore, QUEEN could be useful to estimate the overall homogeneity of the qs within a new family. Nevertheless, as is also true for many other deep learning applications, QUEEN is not able to accurately predict qs changes that occur from point mutation (data not shown). Of note, QUEEN was trained on sequences extracted from solved structure rather than the full protein sequence as reported in uniprot (31) (for which the qs is not necessarily the same and known), and therefore it remains to be investigated how well it will perform for full sequences. In its current implementation, we suggest using it for the prediction of selected regions that mainly include defined domains, as QUEEN did not have the opportunity to train on unstructured regions that are not resolved in solved structures.

A number of qs prediction and assessment tools are already available. The widely used PISA strictly relies on a solved structure - not only the monomeric fold, but the entire complex. In this context, PISA calculates and determines the qs from within the available options in the complex. When a crystal structure is examined by PISA, only the combinations comprising the crystal lattice and the consequent complex are considered. This reduces, often dramatically, the qs that may be considered, improving predictions. Nonetheless, QUEEN succeeds in certain cases where PISA fails (Figure 9 B and C). Pending our ability to reliably identify these cases, this could pave the way for further improvement by a pLM-based model such as QUEEN. Optimal extended leverage of structure prediction can be obtained by using state of the art structure prediction methods such as AlphaFold and ESMfold to provide structural information (e.g. (2)), in particular by modeling different qs states to identify the most promising one (32).

## Conclusions

Protein sequences hold information regarding the protein’s quaternary structure. Leveraging this information can be directed towards predicting the qs from sequence alone, as shown here in detail. This work offers a useful tool for qs prediction, and opens the door to further investigation and improvement of utilizing pLMs to predict qs.

## Methods

### Dataset

The data used in this study is a subset of the data annotated in QSbio V6_2020, courtesy of Emmanuel Levy (16). The data was filtered so that the remaining set met the following criteria: homomers (*i.e*. comprised of copies of the same monomer), with qs annotation (that was not changed in an internal validation process, *i.e*., “corrected_nsub” == “nsub”), and highest confidence level (best biological unit “Best_BU” == 1 and QSBIO_err_prob < 15). Duplicate fasta sequences were removed, retaining only the highest confidence annotation. This process resulted in a final set of 31,994 entries, spanning 19 qs labels. The sequences were taken from the solved PDB structure, including all residues present in the experimental construct.

### Generating independent train and hold-out sets

This filtered set above was split into a train and a hold-out set (using the *sklearn* function *model_selection.StratifiedGroupKFold* (33). To prevent leakage between the data used for training and the used for final validation of the model (the hold-out set), we grouped the sequences by similarity (using MMseqs (34)), and clustered the data by 30% sequence identity with at least 30% coverage:

*mmseqs easy-cluster <input_fasta.fasta> session <session_dir_location> --min-seq-id 0.3 -c 0.3 -s 8 --max-seqs 1000 --cluster-mode 1 --cluster-reassign*.

This relatively low coverage cut-off was selected to make sure the groups are indeed as separated as possible. Grouping of the data was done with the cluster representatives, defining the sets so that an entire cluster is included in the same set. We stratified the sets to ensure similar distributions of qs states in both sets, as much as possible.

About 10% of the data was put aside as a hold-out set. Within the remaining set we performed several cross-validation (cv) experiments for hyper-parameter tuning, model selection etc. For this, we divided this set into 5 sets of approximately similar size, and 5 cv experiments were performed using each time a 80%-20% separation of training and test set (the approximation is due to the group and stratification constraints).

### Generation of embeddings and pooling

Embeddings from the ESM2 model were generated locally, with no need for a GPU. The ESM2 model generates a vector of length 1280 per each amino acid in each protein sequence, resulting in a matrix of size *L×1280* (L = sequence length). We performed mean pooling, averaging the matrix over the L dimension, to obtain a vector of 1280 for each sequence. This results in uniform embedding length for each entry.

Protbert_BFD embeddings were generated similarly, where the representation for each residue is a vector of length 1024, thus resulting in a vector representation of length 1024 per entry.

### Dimensionality reduction

We performed supervised dimensionality reduction with *UMAP* (35). The parameters used are:

*n_components=3, n_neighbors=350, min_dist=0.5*.

After obtaining the reduced vectors we plotted all three target dimensions for visualization.

### Nearest neighbor annotation

Comparison of the embeddings to the sequence was carried out by using the nearest neighbor algorithm to transfer the qs annotations. Annotation transfer was conducted in two parallel implementations: (1) without inclusion of homolog sequences for comparison and annotation transfer *(i.e*., sequences in the test set were assigned a qs based on the closest entry in the train set). (2) including all close homolog sequences as well *(i.e*. including sequences from the test set). The nearest neighbor was identified based on sequence similarity distance calculated with a local alignment as incorporated in MMseqs (34), with default parameters (sequence-based annotation transfer), or based on cosine similarity of the embedding vectors (for pLM-based annotation transfer).

### Hyperparameter tuning and MLP model

Exploring hyperparameters was done using *RandomizedSearchCV* from *sklearn* (33). Among the hyperparameters used was the downsampling of the monomer and the dimer classes, which improved performance and was thus included in the pipeline. For downsampling we used the package *imblearn* and the function *RandomUnderSampler* (36). The final downsampling factor chosen was 3, thus reducing to 33% the amount of monomers and dimers in the training set.

The hyperparameter search was performed on the 5 fold cv described above, and the final parameters were selected based on the best results for adjusted balanced accuracy.

- Model parameters: *MLPClassifier*

> *(activation=‘identity’, alpha=0.0005, batch_size=250, hidden_layer_sizes=(120,), learning_rate= ‘adaptive’, solver= ‘adam’, learning_rate_init=0.01, max_iter= 1000, n_iter_no_change=20, random_state=22, tol=0.001)*
- Sampled space (in bold: chosen parameter value):: *activation = [**‘identity’**, ‘logistic’, ‘tanh’, ‘relu’]* *learning_rate = [‘constant’, ‘invscaling’, **‘adaptive’**]* *learning_rate_init = [0.001,0.005, **0.01**, 0.05, 0.1]* *solver = [‘sgd’, **‘adam’**]* *max_iter = [10, 100, 400, 800, **1000**, 1500]* *n_iter_no_change = [5, 10, 15, **20**, 30, 40]* *tol = [0.0001, 0.0005, **0.001**, 0.01, 0.1, 1]* *hidden_layer_sizes = [(10,), (20,), (40,), (60,), (80,), (100,), **(120,)**, (140,), (160,), (180,), (200,), (40,40,40), (12, 24, 36), (40, 40, 40), (40, 40, 40, 40, 40, 40, 40, 40, 40, 40), (12, 24, 36, 48), (24, 36, 48), (24, 36, 48, 64), (36, 48, 60, 72, 84, 96), (1000,), (2000,)]* *alpha = [0.0001, **0.0005**, 0.001, 0.01, 0.1, 1]* *batch_size= [10, 50, 100, 150, 200, **250**, 400, 600, 1000, 1500, 2000, 5000]*

### Statistical analysis

The statistical analysis was carried out using a one-sided (“greater”) Wilcoxon test, implemented through *scipy.stats.ranksums*, with *alternative=‘greater’*, the correct predictions passed as x and the wrong predictions passed as y.

### Packages and versions

All the scripts were written using *Python3.7*.

Embedding the sequences was carried out using *Torch*. The ESM2 model used is EsmForSequenceClassification, with the specific version of *esm2_t6_8M_UR50D* (23). Additionally was initialized the parameter problem_type=“multi_label_classification”.

Embedding with protbert was done using BertModel and the specific version *prot_bert_bfd* (22). *MMseqs* version ad5837b3444728411e6c90f8c6ba9370f665c443 was used, installed locally. *UMAP* version 0.5 was used.

For analyses and data handling we used *pandas* version 1.3.5 (37).

Graphs and plots were generated with *matplotlib* and *seaborn*, versions 3.5.3 and 0.12.0 (38,39).

The network plot was generated using *networkx* version 3.0.

### Pymol

Protein structure visualization was done using *PyMol* version 2.2.0 Open-Source.

## Declarations

### Ethics approval and consent to participate

Not applicable

### Consent for publication

Not applicable

### Availability of data and materials

all data is provided as supplementary tables in csv format. The data is also available in the github repository: https://github.com/Furman-Lab/QUEEN

### Authors’ contributions

OA and TT conceived the idea for the research. TT performed proof of concepts experiments, and OA performed the experiments for the model presented here. OA, TT and OSF refined the concept and realized the final project and analysis. ZBA and LT assisted with analyses and ideas. OA and OSF wrote the manuscript. OSF supervised the project, OA, TT and OSF acquired funding

## Acknowledgements

The authors thank Prof. Emmanuel Levy for the updated QSbio data and for insightful discussions. The authors, and specifically OA, would like to thank Matan Avraham for fruitful discussions and thoughts, and for his great help with Figure 2C and 7.

## Supplementary Material

### Supplementary Figures

**Supplementary Figure S1:**
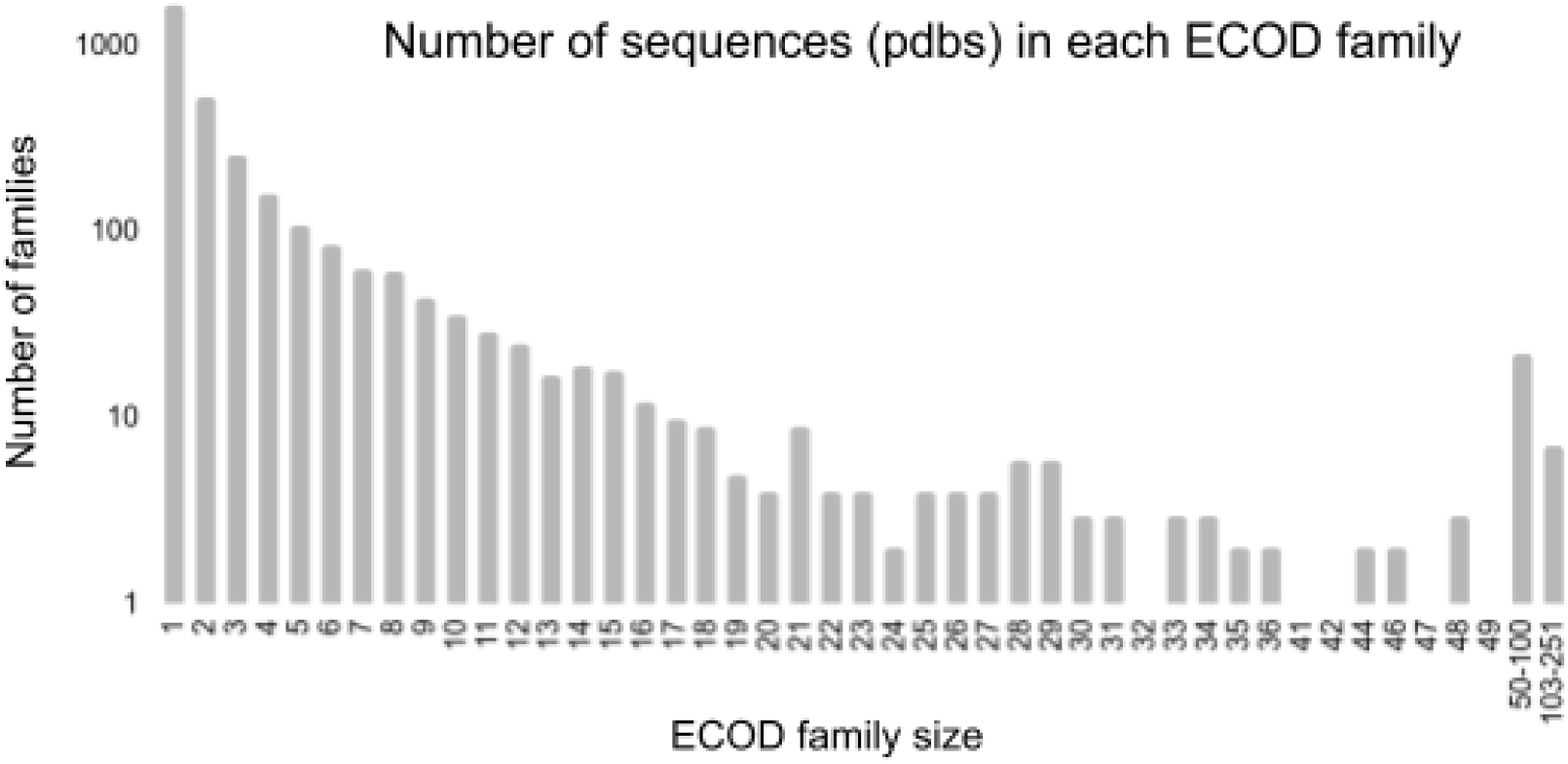
ECOD family sizes. X-axis shows the family size, y-axis the amount of families of the corresponding size. Large families are grouped together for clarity. Accompanies Figure 2.

**Supplementary Figure S2:**
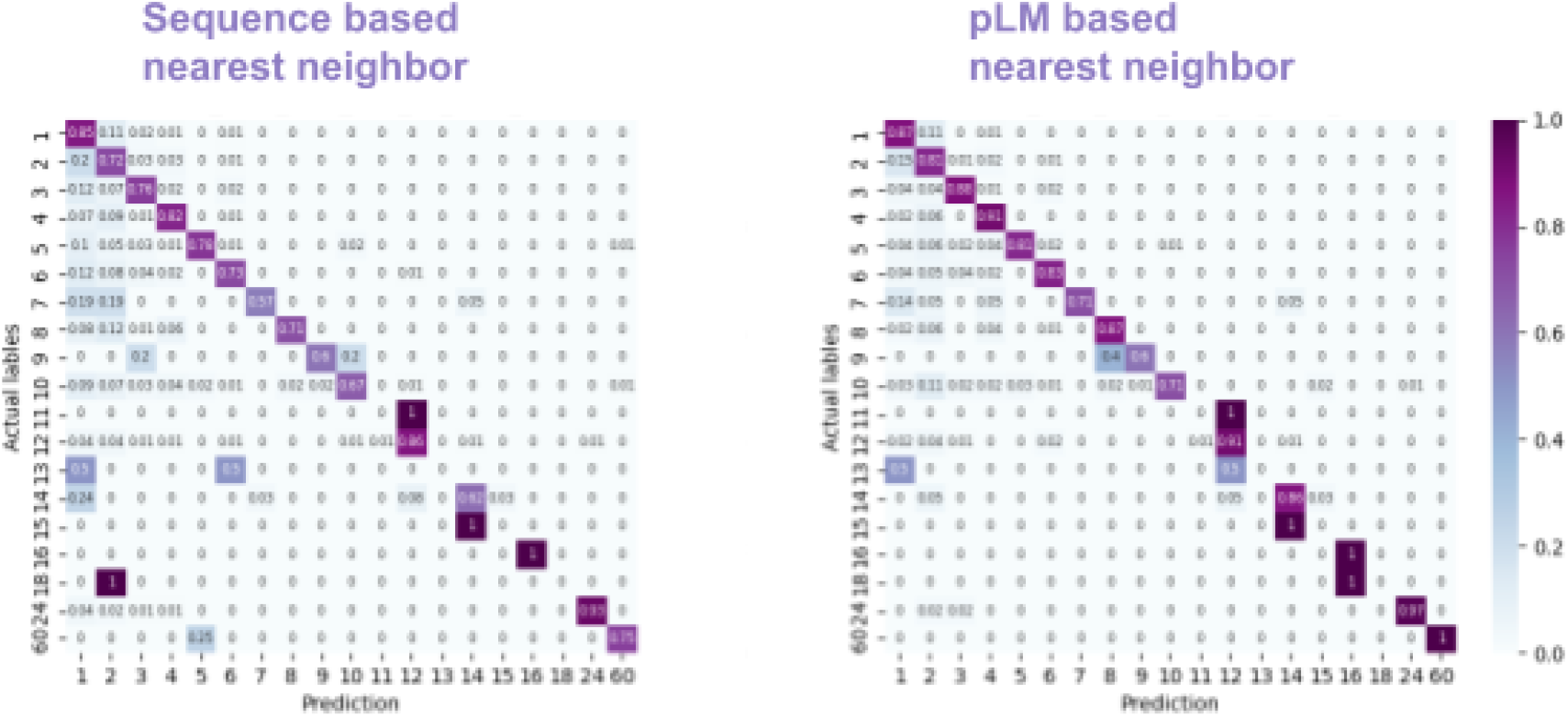
Success of nearest neighbor annotation transfer when close homologs are available. **A**. Sequence-based annotation transfer (balanced accuracy, BA=0.6), **B.** Cosine similarity-based annotation transfer (BA=0.67). Accompanies Figures 4A and B, respectively.

**Supplementary Figure S3:**
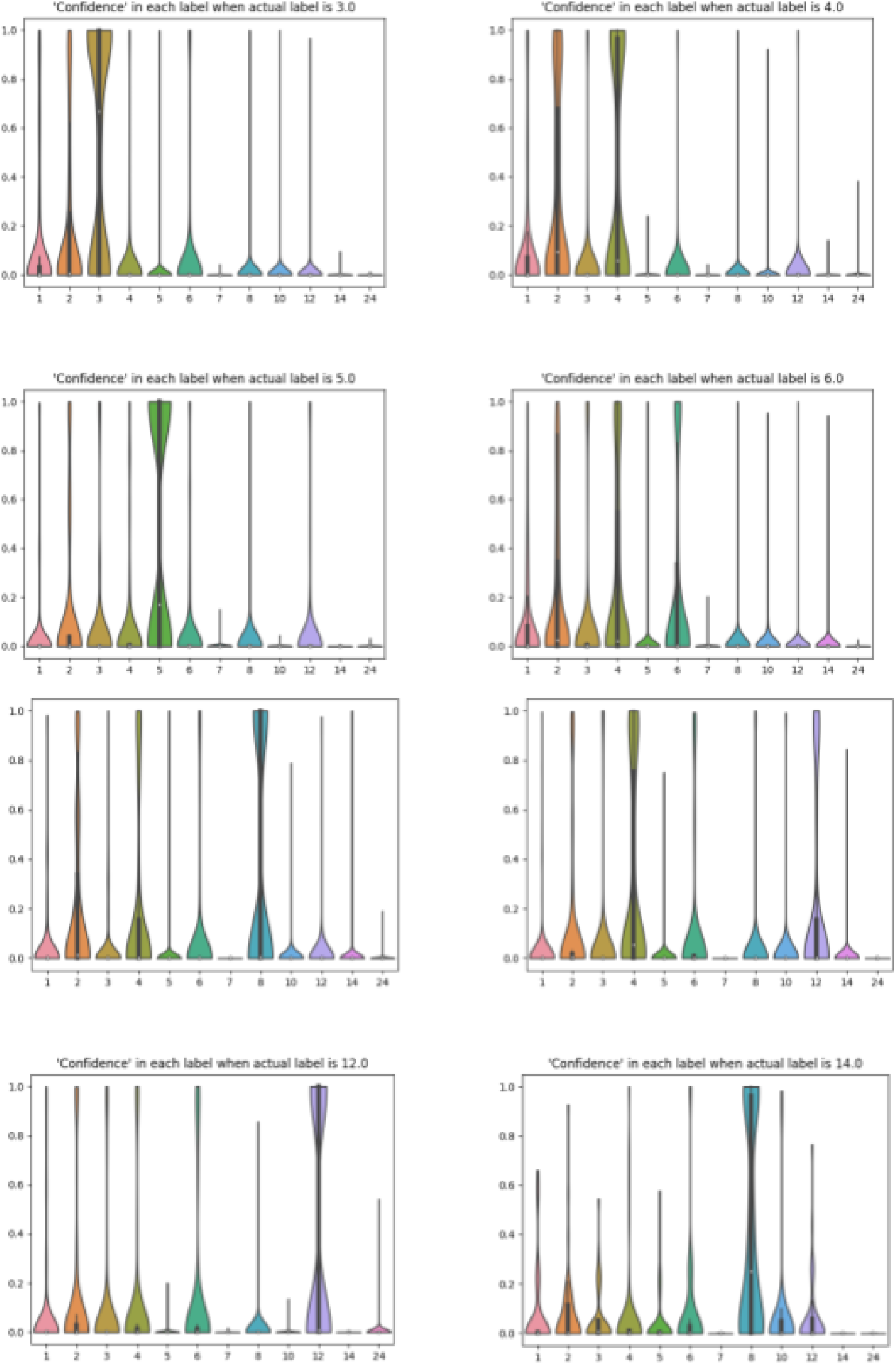
Distribution of predicted probability (‘confidence’} for representative qs categories. The plots show violin plots of ttie prediction probabilities {yoaxis}for different qs labels (x-a»s), for actual qs=3,4,5,6,8,10,12 and 14 (indicated in tha title and highlighted in cyan on the x-axis). Accompanies Figure 7.

**Supplementary Figure S4:**
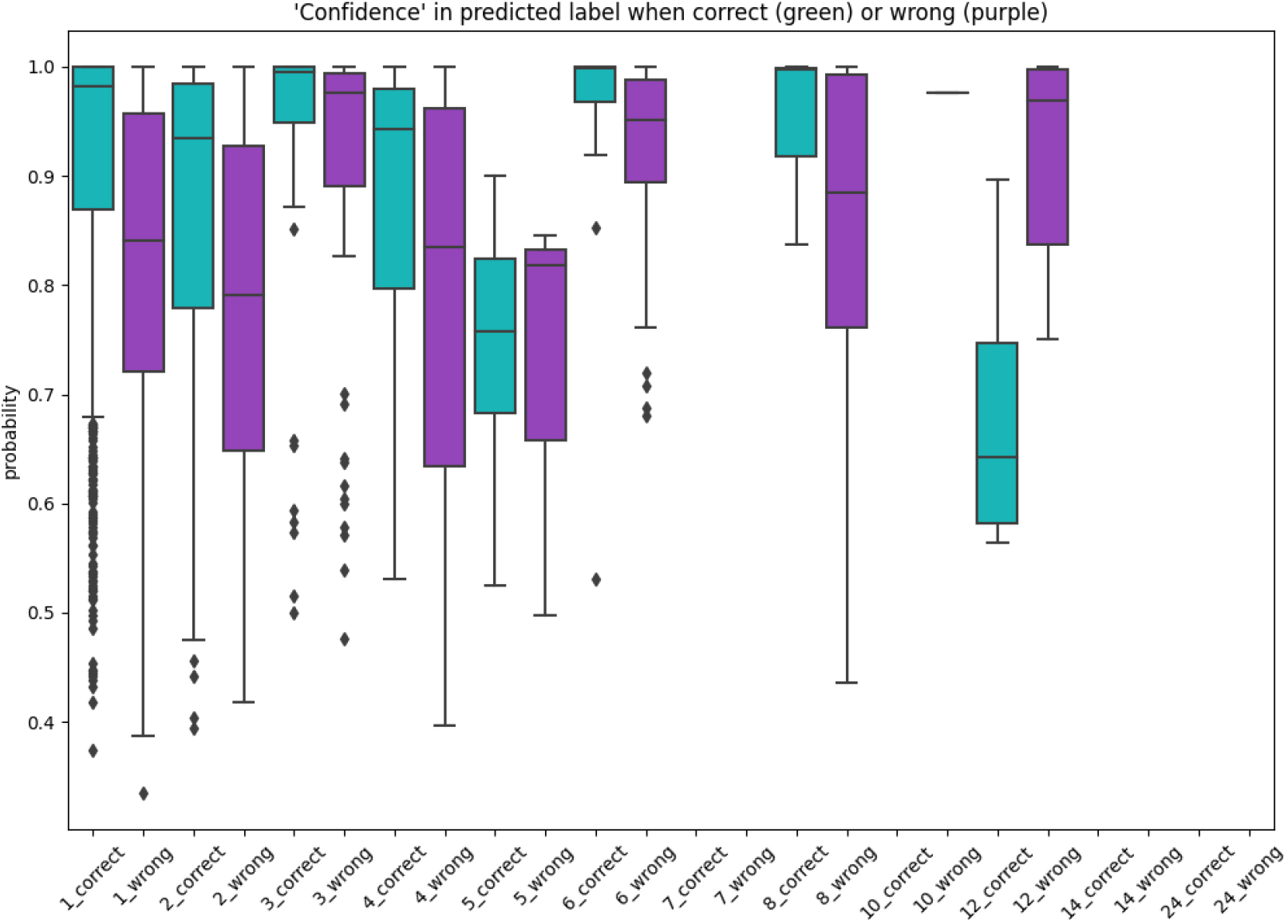
QUEEN predictions shows higher confidence for correct assignments also for the hold out set. Accompanies Figure 10. For each qs (x-axis) box plots and dots show the distribution of the probabilities of correct (green) and incorrect (purple) predictions. Correct predictions are consistently predicted with higher probability than incorrect predictions, for most qs. This separation can be used to assess correct predictions, also in the absence of homologs.

### Supplementary Tables

Supplementary Tables SI and II are attached as tsv files.

**Supplementary Table SIII(a).** Wilcoxon statistics and P-values of full homology annotation transfer

|  |  |  |
| --- | --- | --- |
| qs | statistic | p-value |
| 1 | 38.66 | 0 |
| 2 | 35.44 | 2.05E-275 |
| 3 | 14.40 | 2.46E-47 |
| 4 | 25.92 | 1.96E-148 |
| 5 | 5.02 | 2.61E-07 |
| 6 | 14.14 | 1.10E-45 |
| 7 | 1.64 | 0.051 |
| 8 | 10.10 | 2.69E-24 |
| 9 | 1.96 | 0.025 |
| 10 | 5.00 | 2.89E-07 |
| 11 | nan | nan |
| 12 | 5.82 | 2.95E-09 |
| 13 | nan | nan |
| 14 | 2.93 | 0.002 |
| 15 | nan | nan |
| 16 | 2.24 | 0.013 |
| 18 | nan | nan |
| 24 | 3.74 | 9.10E-05 |
| 60 | nan | nan |

**Supplementary Table SIII(b).** Wilcoxon statistics and P-values of no homology information annotation transfer

|  |  |  |
| --- | --- | --- |
| qs | statistic | p-value |
| 1 | 1.84 | 0.033 |
| 2 | 6.20 | 2.82E-10 |
| 3 | 2.90 | 0.002 |
| 4 | 3.38 | 3.56E-04 |
| 5 | 0.93 | 0.177 |
| 6 | 0.97 | 0.165 |
| 7 | nan | nan |
| 8 | 0.00 | 0.500 |
| 10 | nan | nan |
| 12 | 1.47 | 0.071 |
| 14 | nan | nan |
| 24 | nan | nan |

**Supplementary Table SIV.** Wilcoxon statistics and P-values of QUEENI

| Supplementary Table SIV - Wilcoxon statistics and P-values of QUEENI |  |  |
| --- | --- | --- |
| qs | statistic | p-value |
| 1 | 19.84 | 6.30E-88 |
| 2 | 8.83 | 5.40E-19 |
| 3 | 11.63 | 1.46E-31 |
| 4 | 12.21 | 1.35E-34 |
| 5 | 4.87 | 5.71E-07 |
| 6 | 4.84 | 6.53E-07 |
| 7 | nan | nan |
| 8 | 7.36 | 9.50E-14 |
| 10 | 2.59 | 0.005 |
| 12 | 8.85 | 4.33E-19 |
| 14 | nan | nan |
| 24 | 1.75 | 0.040 |

